# *Paenibacillus glycinicola* sp. nov., *Paenibacillus triticicola* sp. nov., *Pseudescherichia oryzae* sp. nov., *Pseudomonas gossypii* sp. nov., and *Tardiphaga zeae* sp. nov., five novel bacterial species from agricultural crops

**DOI:** 10.1101/2022.11.02.514884

**Authors:** Sarah Seaton, Emily Auker, Allison Connolly, Mekdem Tesfaye, Patrik Inderbitzin, Stephen J. Barnett, Christopher M. M. Franco, Martha E. Trujillo

## Abstract

Five novel species from healthy plants and other substrates are described. These are *Paenibacillus glycinicola* sp. nov. from *Glycine max* in Iowa, United States; *Paenibacillus triticicola* sp. nov. from *Triticum aestivum* in Australia, *Allium cepa* in Texas, garden soil in Wisconsin, cow’s milk in New York state, *Panicum virgatum* in Michigan, sweet sorghum in Nebraska, and *Zea mays* in Ohio; *Pseudescherichia oryzae* sp. nov. from *Oryza sativa* in Arkansas; *Pseudomonas gossypii* sp. nov. from a healthy *Bos taurus* in Kansas, *Capsicum annuum* from Arizona, *Gossypium hirsutum* in New Mexico, Oklahoma, and Texas, and *Zea mays* from Arkansas; and *Tardiphaga zeae* sp. nov. from *Zea mays* in Iowa. No pathogenic strains are known for any of the novel species.

## INTRODUCTION

Agricultural crops have diverse microbiomes (Favela et al. 2022; Kawasaki et al. 2021; Kuźniar et al. 2020; Longley et al. 2020; Wei et al. 2019) that differ between plant species and compartments (Xiong et al. 2021), and play crucial roles for plant health (Berendsen et al. 2012). A better understanding of the interactions between microbiomes, plants, and the environment, is essential for microbiome manipulation to the benefit of increased sustainability and climate change resilience in agroecosystems (Mariotte et al. 2018; Sergaki et al. 2018).

Here we describe five plant-associated strains, representing five novel species in the genera *Paenibacillus, Pseudescherichia, Pseudomonas*, and *Tardiphaga. Paenibacillus glycinicola* strain SS118 was isolated from healthy field-grown *Glycine max* in Iowa, United States; *Paenibacillus triticicola* strain PI119 was isolated from the roots of *Triticum aestivum* grown in field soil heated to 80°C from South Australia, Australia; *Pseudescherichia oryzae* strain SS120 was isolated from field-grown *Oryza sativa* in Arkansas, United States; *Pseudomonas gossypii* strain PI121 was isolated from healthy field-grown *Gossypium hirsutum* in New Mexico, United States; and *Tardiphaga zeae* strain SS122 was isolated from healthy field-grown *Zea mays* in Iowa, United States. We provide phenotypic and phylogenomic details describing these five species, and report geographic distribution and host range based on comparison to whole genome assemblies from GenBank of conspecific, cultured strains.

## METHODS

### Isolation

Strain SS118 was isolated from a stem of healthy field-grown *Glycine max* in Iowa, United States; strain PI119 was isolated from the roots of *Triticum aestivum* grown in field soil heated to 80°C, the soil was collected in South Australia, Australia; strain SS120 was isolated from a field-grown *Oryza sativa* seedling in Arkansas, United States; strain PI121 was isolated from a cotton bowl of a healthy field-grown *Gossypium hirsutum* in New Mexico, United States; and strain SS122 was isolated from a root of healthy field-grown *Zea mays* in Iowa, United States. For strains PI121, SS118, SS120, and SS122, plant tissue was washed with a mild detergent to remove particulates, surface-sterilized with bleach (1% v/v sodium hypochlorite) and ethanol (70% v/v), and homogenized. Serial dilutions of tissue homogenate were plated on a panel of media types for endophyte cultivation. For strain PI119, the roots were washed well, heated to 80°C for 30 min, and five-millimeter segments were placed onto Baker’s Yeast agar (BYA1: 4g/L baker’s yeast, 10 g/L agar, 100 mg/L cycloheximide) incubated at 25°C. All strains were streaked to purity and stored in glycerol (20% v/v) at -80°C until subjected to further testing.

### Motility

The strains were tested for flagellar-dependent swimming and swarming motility on R2A plates solidified with 0.3% and 0.6% agar, respectively. Three independent colonies were inoculated onto R2A broth and grown for 36 hr at 24°C. Broth cultures were normalized to an OD600 of 0.1, and 1.5 µl of culture was spotted directly onto the surface of the motility agar. The diameter of colony expansion was measured for 5 days.

### Carbon source utilization

Substrate utilization was assessed using Biolog GenIII Microplates (Catalogue No. 1030) (Biolog Inc., Hayward, CA). Each bacterium was inoculated in duplicate plates using Protocol A, described by the manufacturer, except that plates were incubated at 30°C for 24-48 hours, and *Tardiphaga zeae* strain SS122 required 8 days of incubation. Respiration leading to reduction of the tetrazolium indicator was measured by absorbance at 590 nm.

### Biochemical analyses

Catalase activity was evaluated by immediate effervescence after applying 3% (v/v) hydrogen peroxide solution via the tube method; a positive reaction was indicated by the production of bubbles. *Staphylococcus aureus* NCIMB 12702 and *Streptococcus pyogenes* ATCC 19615 were used as positive and negative controls, respectively. Oxidase activity was evaluated via the oxidation of Kovács oxidase reagent, 1% (w/v) tetra-methyl-p-phenylenediamine dihydrochloride in water, via the filter-paper spot method. A positive reaction was indicated when the color of the microorganism changed to dark purple. *Pseudomonas aeruginosa* NCIMB 12469 and *Escherichia coli* ATCC 25922 were used as positive and negative controls, respectively. Urease activity was evaluated via the hydrolysis of urea in Christensen’s Urea Agar, using phenol red as a pH indicator. *Proteus hauseri* ATCC 13315 and *Escherichia coli* ATCC 25922 were used as positive and negative controls, respectively. Gram staining was performed using standard protocols.

### Phylogenetic and genomic analyses

DNA was extracted from pure cultures using the Omega Mag-Bind Universal Pathogen Kit according to manufacturer’s protocol with a final elution volume of 60µl (Omega Biotek Inc., Norcross, GA). DNA samples were quantified using a Qubit fluorometer (ThermoFisher Scientific, Waltham, MA) and normalized to 100 ng. DNA was prepped using a Nextera DNA Flex Library Prep kit according to the manufacturer’s instructions (Illumina Inc., San Diego, CA). DNA libraries were quantified via qPCR using a KAPA Library Quantification kit (Roche Sequencing and Life Science, Wilmington, MA) and combined in equimolar concentrations into one 24-sample pool. Libraries were sequenced on a MiSeq using pair-end reads (2×200bp). Reads were trimmed of adapters and low-quality bases using Cutadapt (version 1.9.1) and assembled into contigs using MEGAHIT (version 1.1.2) (Mitter et al. 2013). Reads were mapped to contigs using Bowtie2 (version 2.3.4) (Langmead and Salzberg 2012), and contigs were assembled into scaffolds using BESST (2.2.8) (Sahlin et al. 2014). Phylogenomic trees were generated using GToTree (version 1.2.1) (Lee 2019).

16S rRNA gene sequences were extracted from genome assemblies using barrnap (Seemann 2019), and 16S rRNA gene phylogenetic analyses were performed using FastTree (Price et al. 2010) using a General Time Reversible substitution model. Taxon sampling for each species is described in the respective phylogenetic tree figure legend.

Average nucleotide identity (ANI) analyses between genome assemblies were performed using fastANI (Jain et al. 2018).

Geographic distribution and host range of novel species were inferred by ANI to assemblies from unidentified species from GenBank (Ciufo et al. 2018) and the Indigo internal collection. An ANI threshold of ≥95% indicated conspecificity (Chun et al. 2018; Richter and Rosselló-Móra 2009).

## RESULTS

### Phylogenetic and genomic analyses

#### *Paenibacillus glycinicola* sp. nov. strain SS118

The strain SS118 16S rRNA gene sequence OP556612 shared 99.6% identity with the 16S of *Paenibacillus azotifigens* NF2-4-5^T^. A phylogenetic tree based on 16S using FastTree (Price et al. 2010) confirmed the affiliation of strain SS118 to the genus *Paenibacillus*. SS118 was most closely related to *Paenibacillus helianthi* P26E ^T^ and *Paenibacillus sonchi* X19-5^T^ with 82% support (Figure 1). The top average nucleotide identity (ANI) was to *Paenibacillus riograndensis* SBR5^T^ with 83.4%, well below the threshold for species demarcation (Chun et al. 2018; Richter and Rosselló-Móra 2009). This showed that strain SS118 represents a new genomic species of *Paenibacillus*.

**Figure 1.**
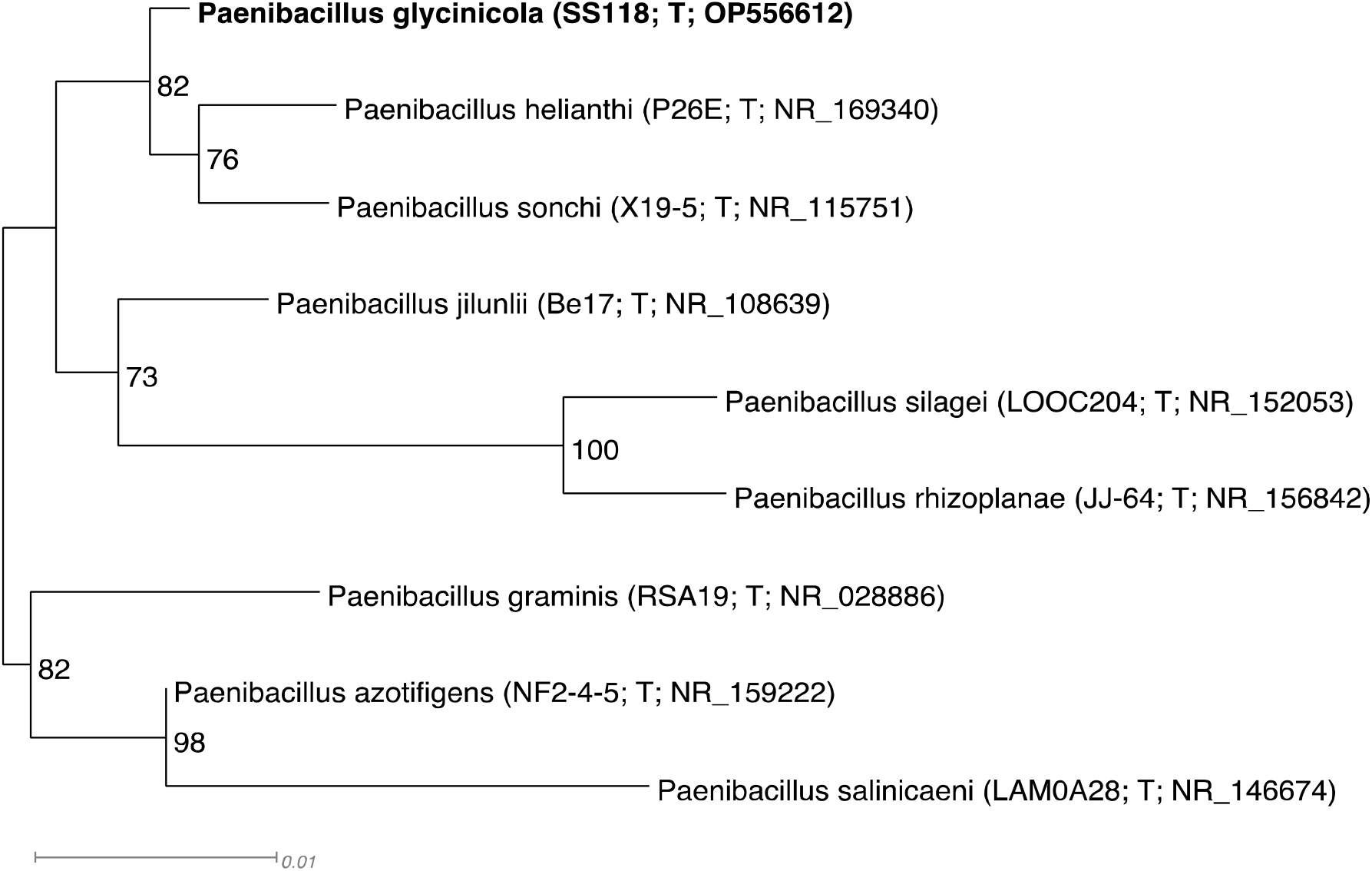
Phylogenetic 16S tree of *Paenibacillus glycinicola* sp. nov. and relatives generated using FastTree (Price et al. 2010). The eight species with the highest 16S identities to *P. glycinicola* were included in the tree, the tree is midpoint rooted. *Paenibacillus glycinicola* is marked in bold. Strain numbers and GenBank accession numbers follow species names, T stands for ‘type’. Support values above 50% are given by the branches. *Paenibacillus glycinicola* is most closely related to *P. helianthi* and *P. sonchi* with 82% support. Branch lengths are proportional to the changes along the branches, a scale bar is provided.

#### *Paenibacillus triticicola* sp. nov. strain PI119

The strain PI119 16S rRNA gene sequence OP556609 shared 99.9% identity with the 16S of *Paenibacillus peoriae* NBRC 15541^T^. A phylogenetic tree based on 16S rRNA gene sequences using FastTree (Price et al. 2010) confirmed the affiliation of strain PI119 to the genus *Paenibacillus*. PI119 was most closely related to *P. peoriae* and *Paenibacillus kribbensis* AM49^T^ with 76% support (Figure 2). The top average nucleotide identity (ANI) was to *Paenibacillus polymyxa* ATCC 842^T^ with 89.8%, well below the threshold for species demarcation of 95% ANI (Chun et al. 2018; Richter and Rosselló-Móra 2009). This showed that strain PI119 represents a new genomic species of *Paenibacillus*.

**Figure 2.**
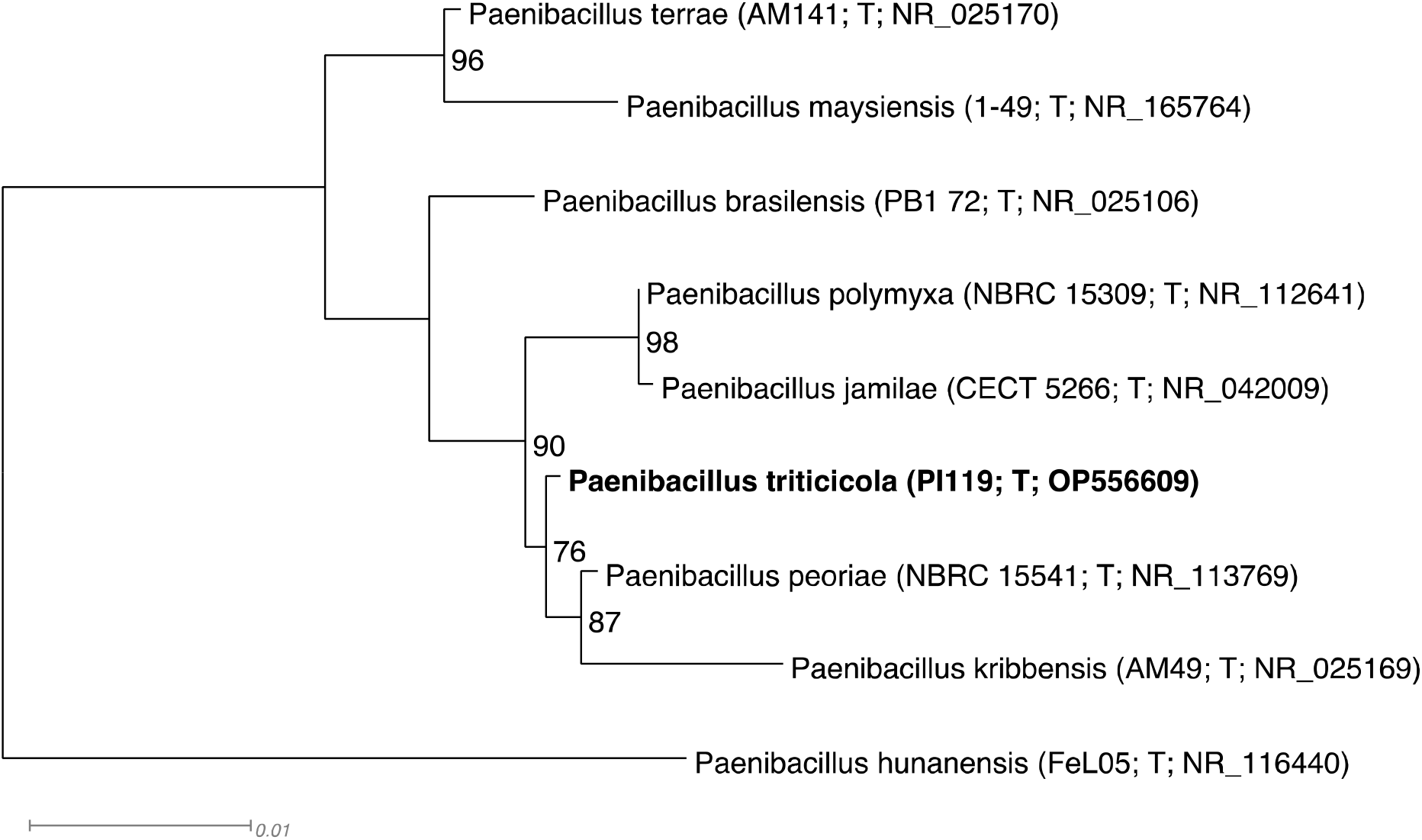
Phylogenetic 16S tree of *Paenibacillus triticicola* sp. nov. and relatives generated using FastTree (Price et al. 2010). The eight species with the highest 16S identities to *Paenibacillus triticicola* were included in the tree, the tree is midpoint rooted. *Paenibacillus triticicola* is marked in bold. Strain numbers and GenBank accession numbers follow species names, T stands for ‘type’. Support values above 50% are given by the branches. *Paenibacillus triticicola* is most closely related to *P. peoriae* and *P. kribbensis* with 76% support. Branch lengths are proportional to the changes along the branches, a scale bar is provided.

#### *Pseudescherichia oryzae* sp. nov. strain SS120

The strain SS120 16S rRNA gene sequence OP556613 shared 98.3% identity with the 16S of *Klebsiella aerogenes* ATCC 13048^T^. Due to this low top 16S identity, and the fact that the 16S is not a reliable phylogenetic marker in the *Enterobacteriaceae* (Dauga 2002), we performed a phylogenomic analysis using GToTree (Lee 2019) with all 261 ex-type assemblies of the *Enterobacteriaceae* in GenBank. Strain SS120 was most closely related to *Pseudescherichia vulneris* NCTC12130^T^ with 100% support (Figure 3). The top average nucleotide identity (ANI) was to *P. vulneris* with 82.8%, well below the threshold for species demarcation (Chun et al. 2018; Richter and Rosselló-Móra 2009). This showed that strain SS120 represents a new genomic species of *Pseudescherichia*.

**Figure 3.**
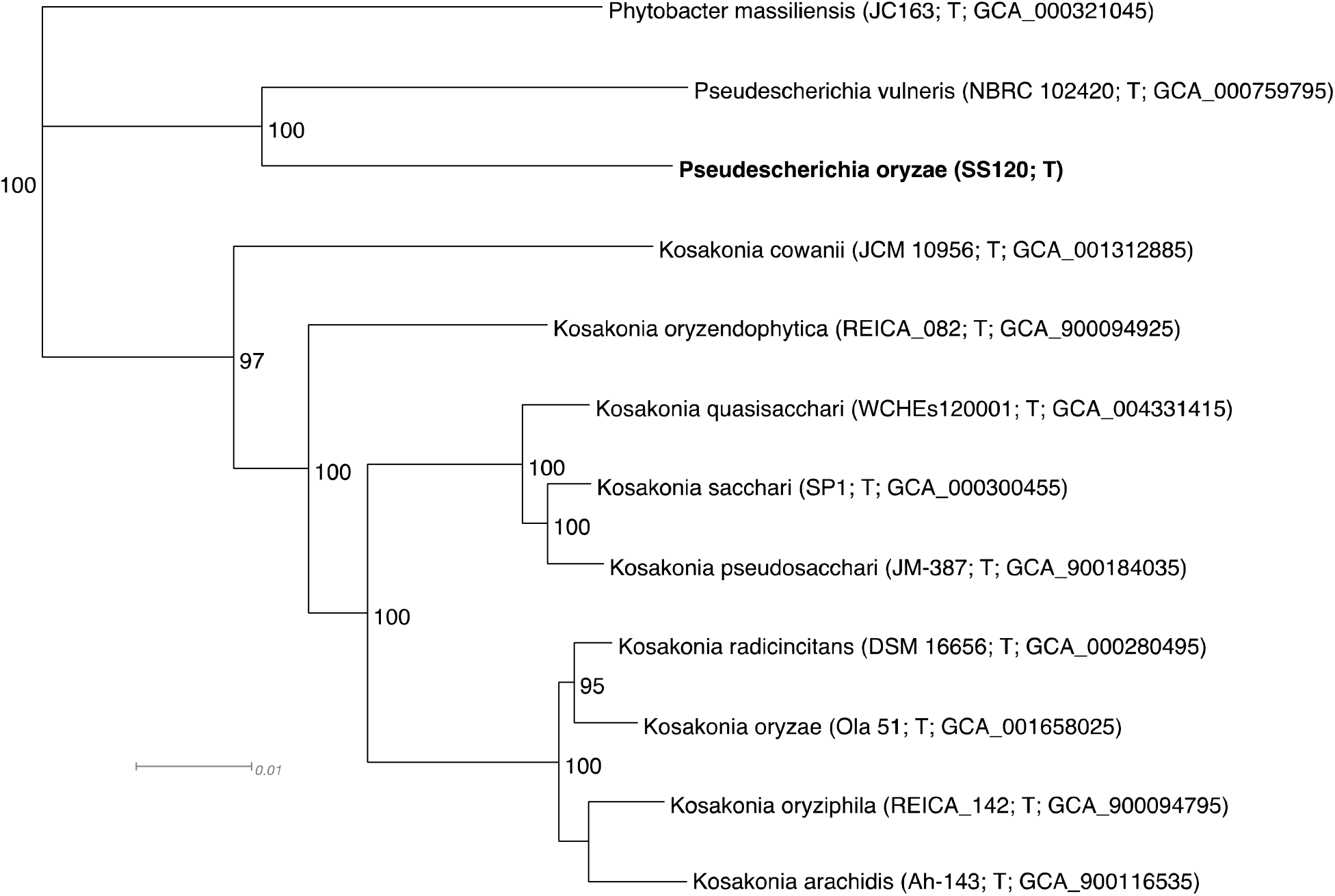
Phylogenomic tree of *Pseudescherichia oryzae* sp. nov. and relatives inferred by GToTree (Lee 2019). Taxon sampling includes the closest relatives of *Pseudescherichia oryzae* based on an *Enterobacteriaceae* species phylogenies (not shown). *Pseudescherichia oryzae* is marked in bold. Strain numbers and GenBank accession numbers follow species names, T stands for ‘type’. Support values above 50% are given by the branches. *Pseudescherichia oryzae* is most closely related to *Pseudescherichia vulneris* with 100% support. Branch lengths are proportional to the changes along the branches, a scale bar is provided.

#### *Pseudomonas gossypii* sp. nov. strain PI121

The strain PI121 16S rRNA gene sequence OP556610 shared 99.3% identity with the 16S of *Pseudomonas oryzihabitans* LMG 7040^T^. A phylogenetic tree using FastTree (Price et al. 2010) confirmed the affiliation of strain PI121 to the genus *Pseudomonas*. PI121 was most closely related to *P. oryzihabitans* and *Pseudomonas oleovorans* IAM 1508^T^ with 100% support (Figure 4). The top average nucleotide identity (ANI) was to *P. oryzihabitans* with 91.2%, well below the threshold for species demarcation (Chun et al. 2018; Richter and Rosselló-Móra 2009), and showing that strain PI121 represents a new genomic species of *Pseudomonas*.

**Figure 4.**
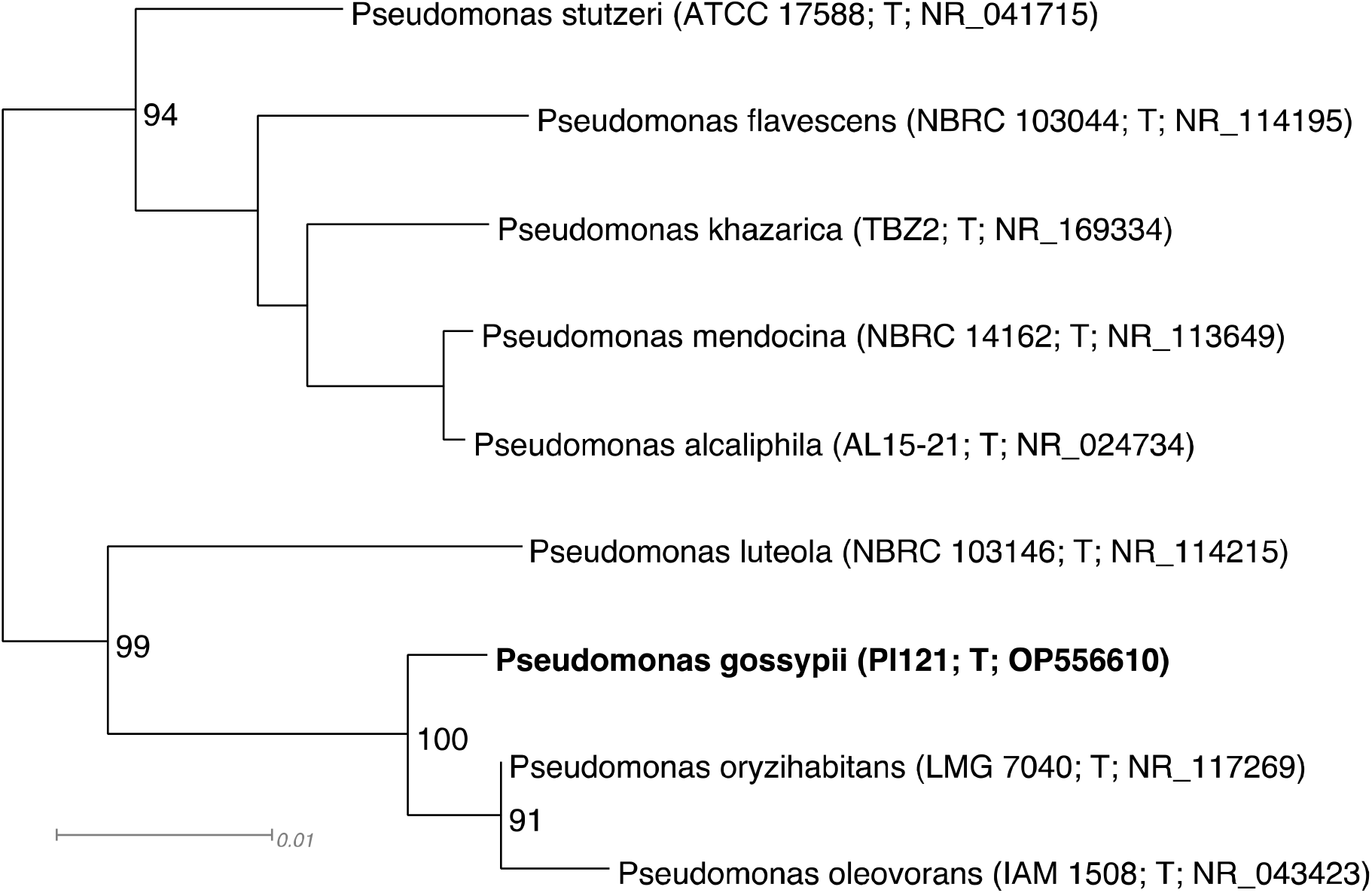
Phylogenetic 16S tree of *Pseudomonas gossypii* sp. nov. and relatives generated using FastTree (Price et al. 2010). The eight species with the highest 16S identities to *Paenibacillus gossypii* were included in the tree, the tree is midpoint rooted. *Pseudomonas gossypii* is marked in bold. Strain numbers and GenBank accession numbers follow species names, T stands for ‘type’. Support values above 50% are given by the branches. *Pseudomonas gossypii* is most closely related to *P. oryzihabitans* and *P. oleovorans* with 100% support. Branch lengths are proportional to the changes along the branches, a scale bar is provided.

#### *Tardiphaga zeae* sp. nov. strain SS122

The strain SS122 16S rRNA gene sequence OP556611 shared 99.5% identity with the 16S rRNA gene sequence of *Tardiphaga robiniae* R-45977^T^. A phylogenetic tree using FastTree (Price et al. 2010) confirmed the affiliation of strain SS122 to the genus *Tardiphaga*. SS122 was most closely related to *T. robiniae* with 100% support (Figure 5). The top average nucleotide identity (ANI) was to *T. robiniae* with 91.8%, well below the threshold for species demarcation (Chun et al. 2018; Richter and Rosselló-Móra 2009), and showing that strain PI121 represents a new genomic species of *Tardiphaga*.

**Figure 5.**
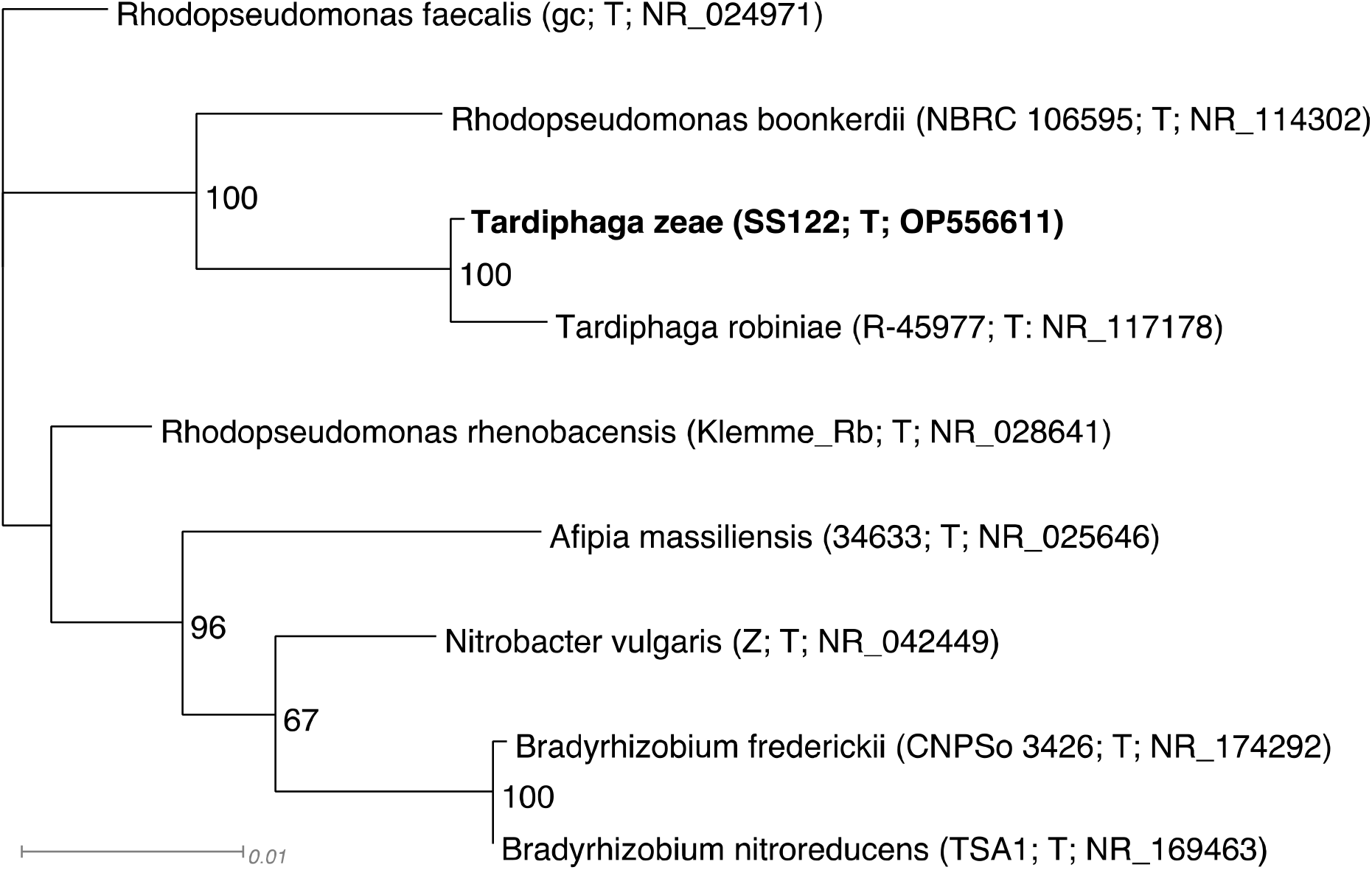
Phylogenetic 16S tree of *Tardiphaga zeae* sp. nov. and relatives generated using FastTree (Price et al. 2010). The eight species with the highest 16S identities to *Tardiphaga zeae* were included in the tree, the tree is midpoint rooted. *Tardiphaga zeae* is marked in bold. Strain numbers and GenBank accession numbers follow species names, T stands for ‘type’. Support values above 50% are given by the branches. *Tardiphaga zeae* is most closely related to *Tardiphaga robiniae* with 100% support. Branch lengths are proportional to the changes along the branches, a scale bar is provided.

### Geographic distribution and host range

Geographic distribution and host range of the novel species was inferred by comparison to congeneric genome assemblies from GenBank and the Indigo internal collection. Distribution maps of the individual species for the United States are included (Figure 6, Figure 7, Figure 8, Figure 9, Figure 10). A summary of the known substrates is provided in Table 1.

**Figure 6.**
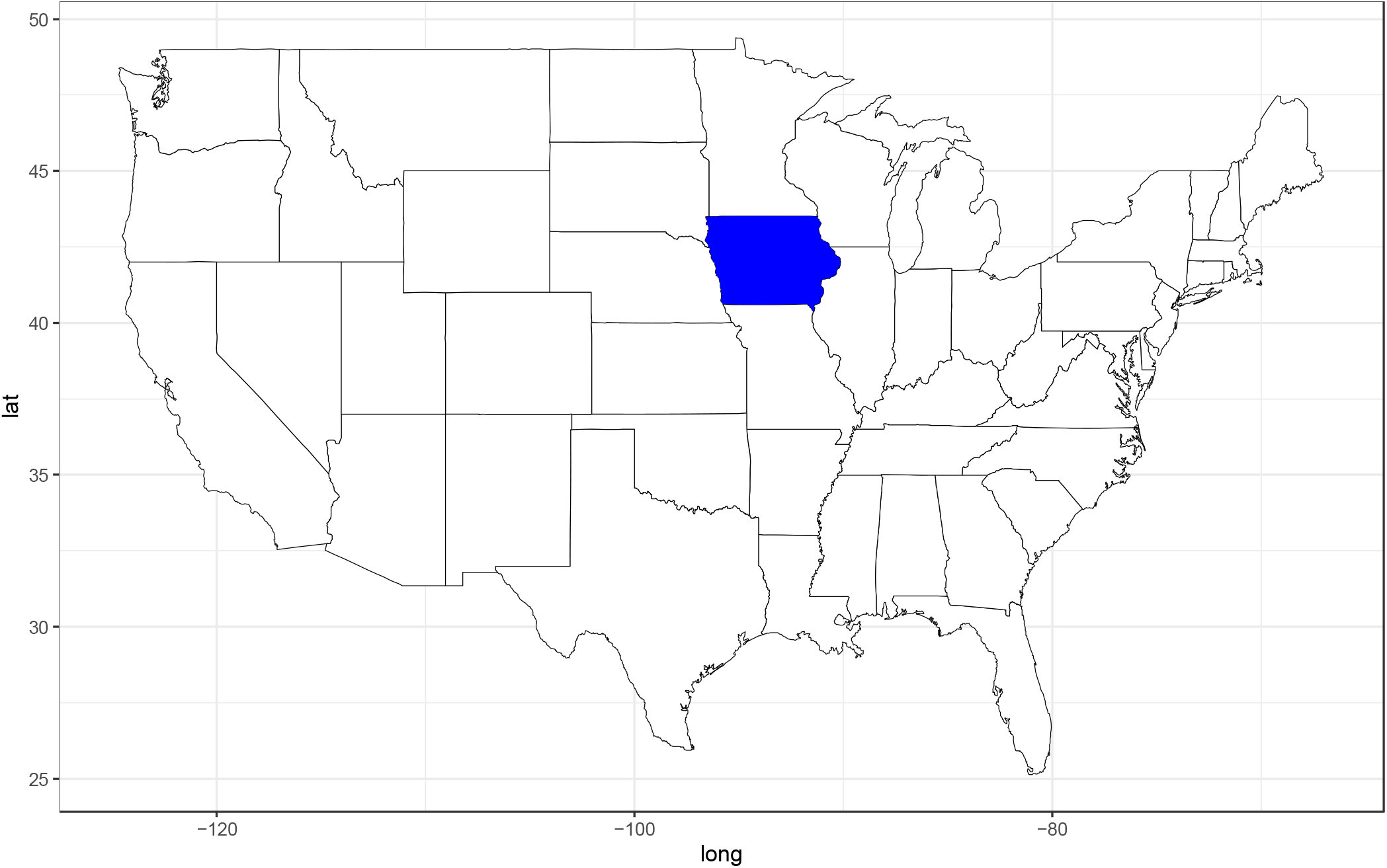
Geographic distribution of *Paenibacillus glycinicola* sp. nov. in the United States from culturing. Dark blue indicates the state of origin of the type strain. See text for details.

**Figure 7.**
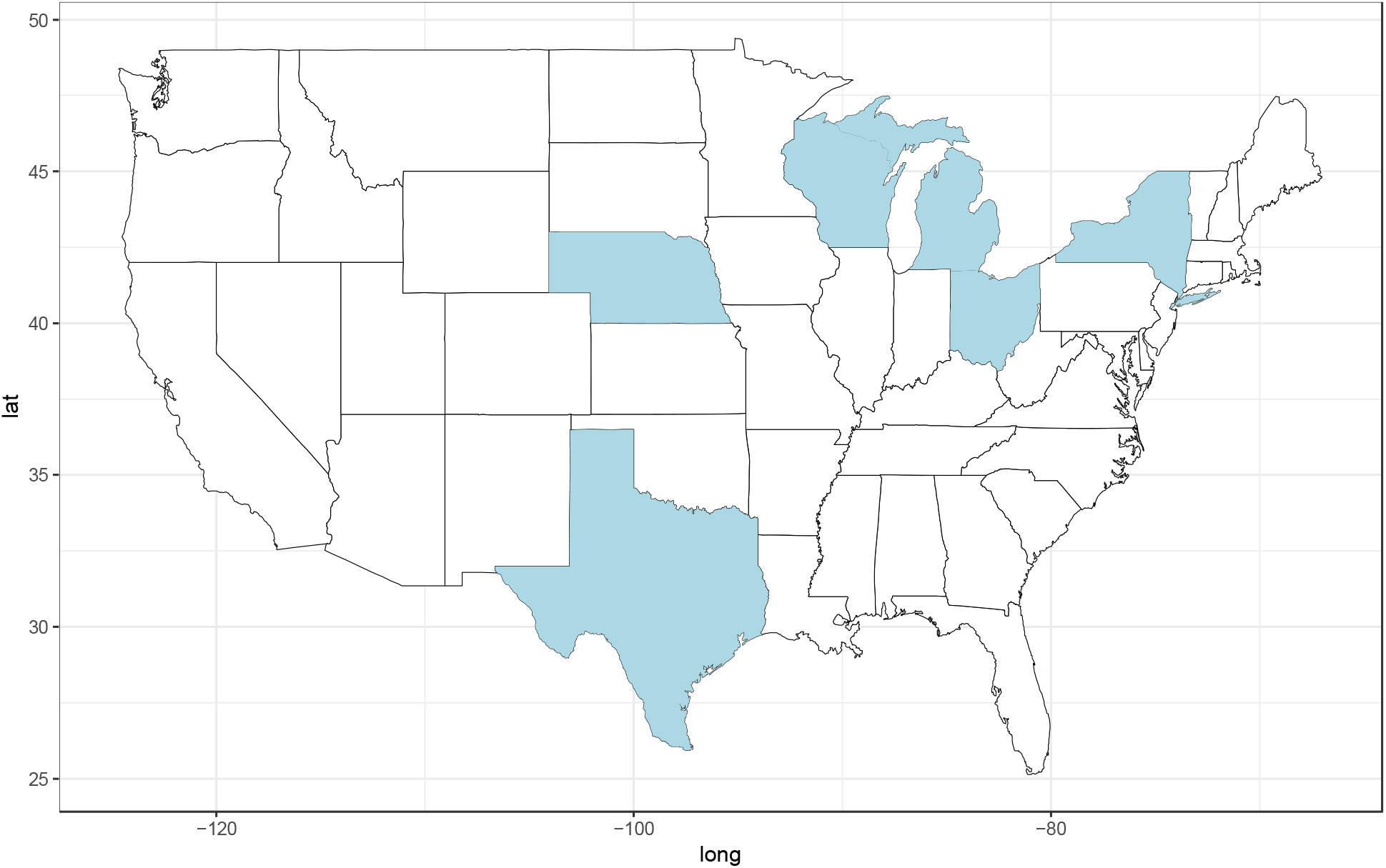
Geographic distribution of *Paenibacillus triticicola* sp. nov. in the United States from culturing. Light blue indicates the state of origin of *Paenibacillus triticicola* strains based on genome assemblies from GenBank and the Indigo internal collection. The type of *Paenibacillus triticicola* was isolated in Australia. See text for details.

**Figure 8.**
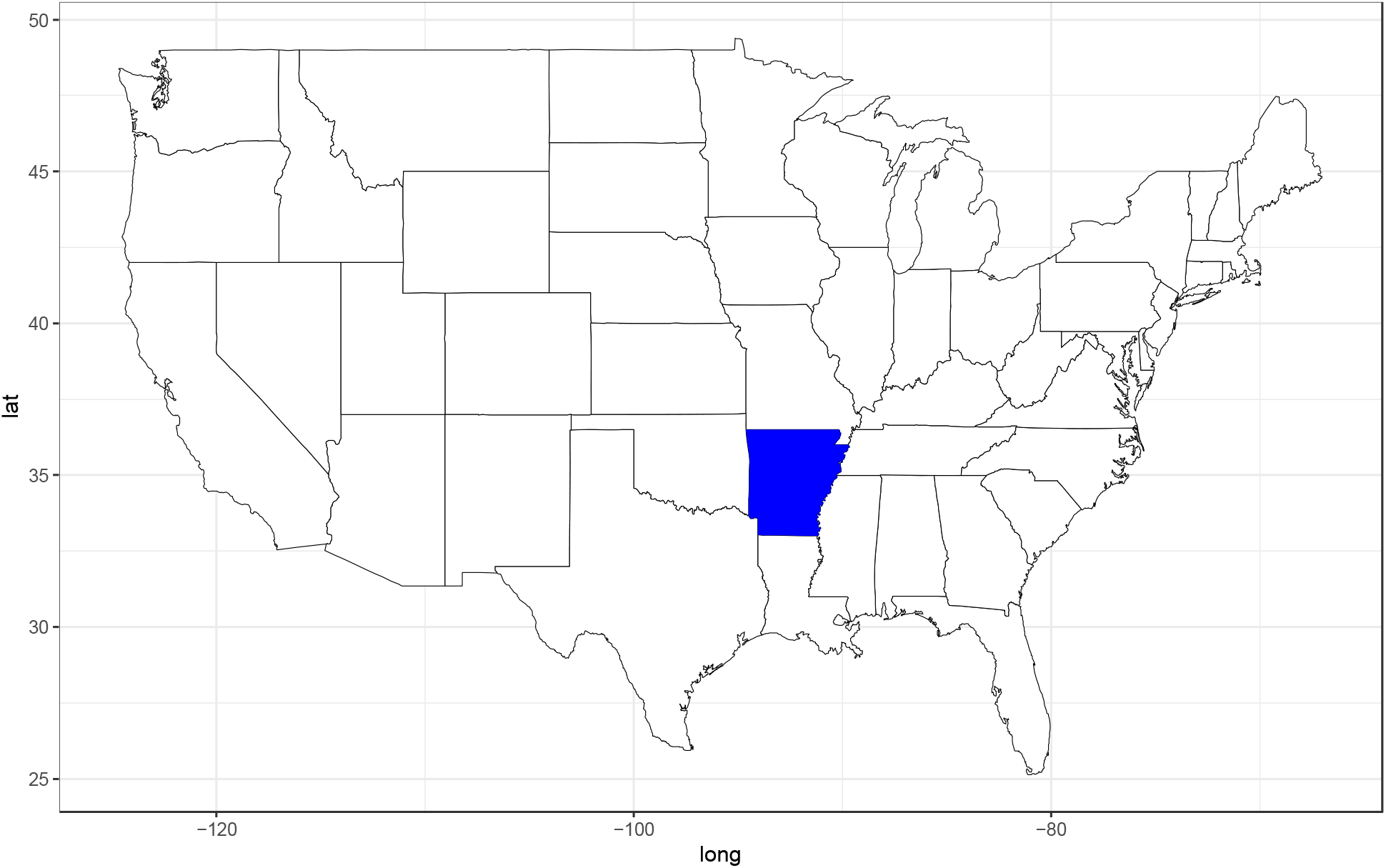
Geographic distribution of *Pseudescherichia oryzae* sp. nov. in the United States from culturing. Dark blue indicates the state of origin of the type strain. See text for details.

**Figure 9.**
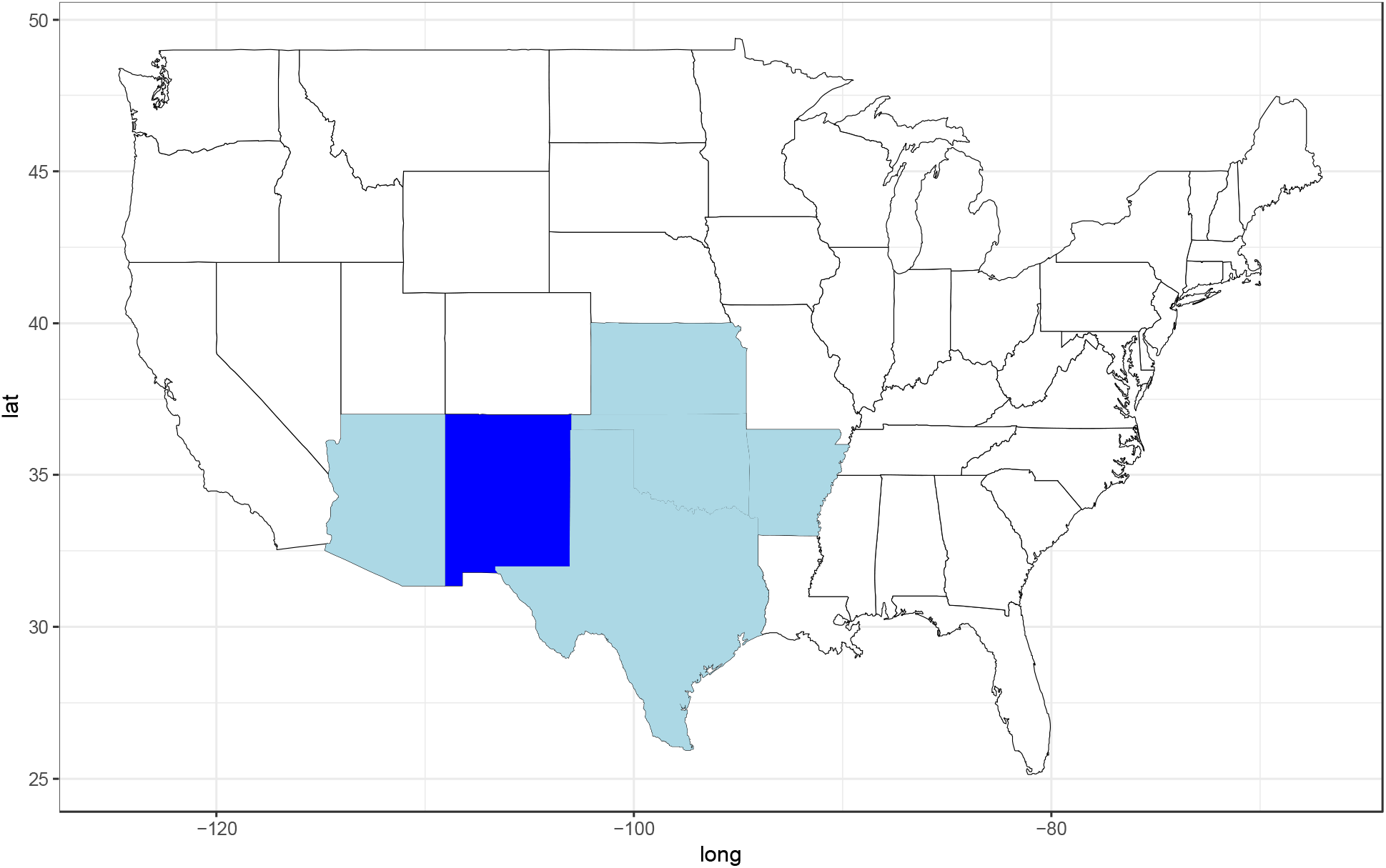
Geographic distribution of *Pseudomonas gossypii* sp. nov. in the United States from culturing. Dark blue indicates the state of origin of the type strain, light blue is the state of origin of additional *Pseudomonas gossypii* strains based on genome assemblies from GenBank and the Indigo internal collection. See text for details.

**Figure 10.**
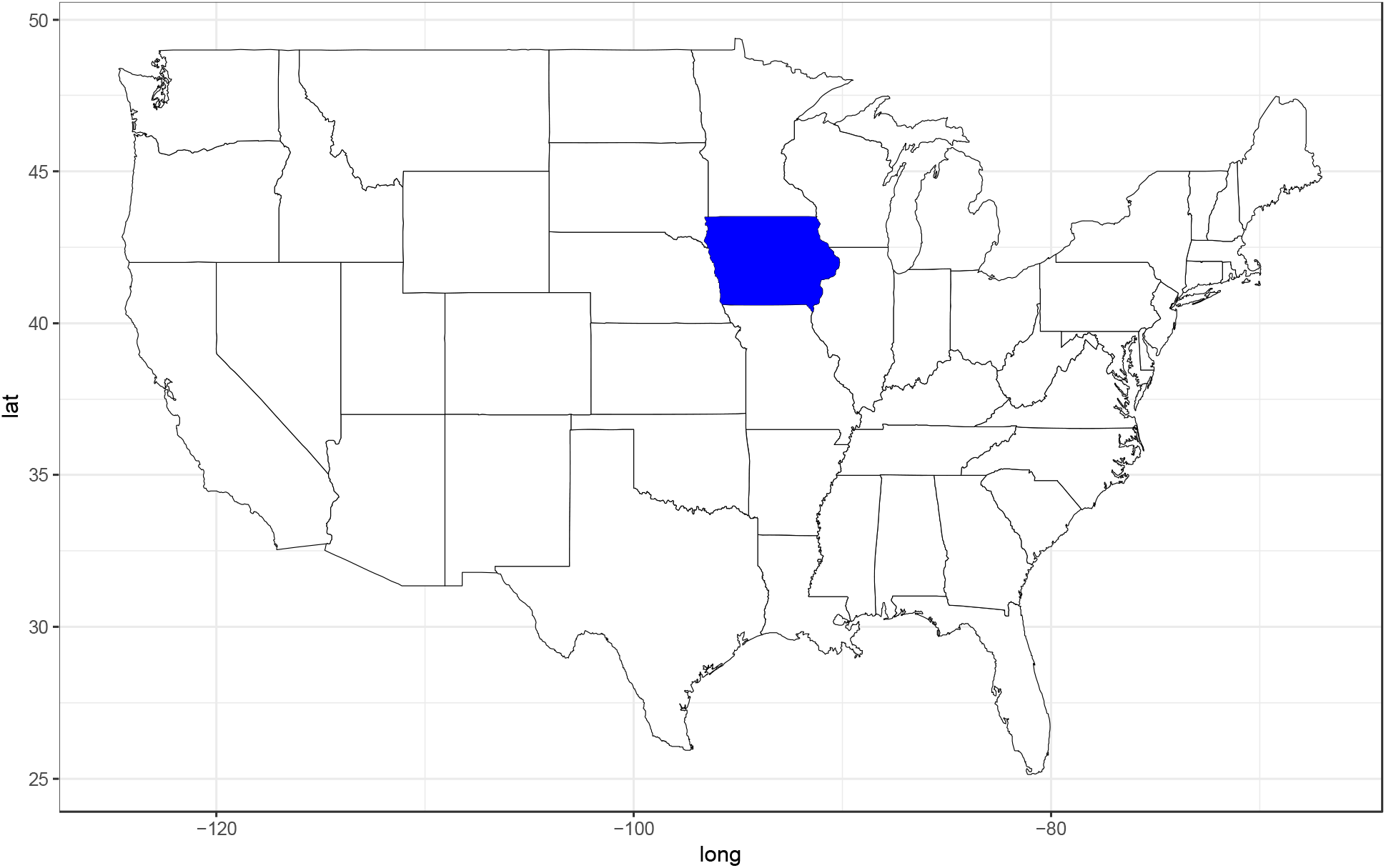
Geographic distribution of *Tardiphaga zeae* sp. nov. in the United States from culturing. Dark blue indicates the state of origin of the type strain. See text for details.

**Table 1.**
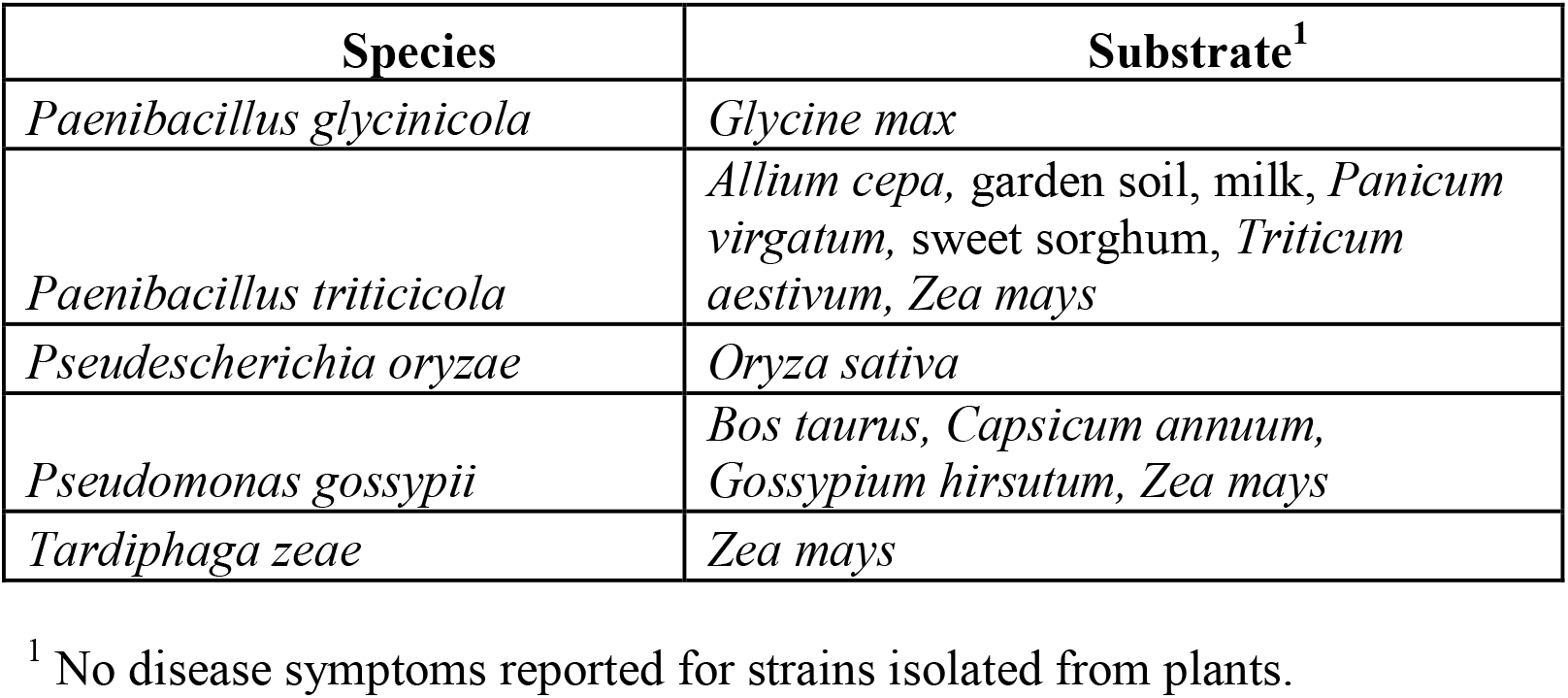
Substrates of novel species based on culturing.

#### *Paenibacillus glycinicola* sp. nov. strain SS118

There were no hits of the strain SS118 genome assembly against GenBank and Indigo genomes at an ANI ≥ 95% threshold. The known geographic distribution of *Paenibacillus glycinicola* sp. nov. in the United States is illustrated in Figure 6.

#### *Paenibacillus triticicola* sp. nov. strain PI119

Hits to strain PI119 included the following GenBank assemblies: Assembly GCF_009827695.1 from the rhizosphere of *Allium cepa* in Texas (Garcia et al. 2020) (ANI: 95.5%; query coverage: 63.7%), assembly GCF_022537985.1 from garden soil in Wisconsin, (ANI: 95.5%, query coverage: 87.0%), five assemblies from milk in New York state (Trmčić et al. 2015) (GCF_001955935.1, GCF_001956115.1, GCF_001956155.1, GCF_001956225.1, GCF_001956235.1; ANI range: 95.5 – 95.9%, query coverage range: 86.7 – 88.3%), assembly GCF_017833155.1 from the rhizosphere of *Panicum virgatum* in Michigan (ANI: 96.0%, query coverage: 87.8%), and assembly GCF_017833915.1 from the roots of sweet sorghum (ANI: 95.1%, query coverage: 86.0%) from Nebraska. From the Indigo internal collection, there was a hit to the assembly of strain P123 from *Zea mays* from Ohio (Figure 7).

#### *Pseudescherichia oryzae* sp. nov. strain SS120

There were no hits of the strain SS120 genome assembly against GenBank and Indigo genomes at an ANI ≥ 95% threshold. The known geographic distribution of *Pseudescherichia oryzae* sp. nov. in the United States is illustrated in Figure 8.

#### *Pseudomonas gossypii* sp. nov. strain PI121

There was one hit among GenBank assemblies to strain PI121, which was GCF_001518815.1 from the nasopharynx of a healthy calf (*Bos taurus*) in Kansas. Among the Indigo collection, there were hits against strain SS124 from a leaf of *Capsicum annum* in Arizona (ANI: 98.2%, query coverage: 91.2%), strain PI125 from a leaf of *Gossypium hirsutum* in Oklahoma (ANI: 98.0%, query coverage: 94.3%), strain SS126 from a seedling of *Gossypium hirsutum* in Texas (ANI: 98.1%, query coverage: 91.2%), and strain PI127 from a seedling of *Zea mays* from Arkansas (ANI: 98.0%, query coverage: 90.9%) (Figure 9).

#### *Tardiphaga zeae* sp. nov. strain SS122

There were no hits of the strain SS122 genome assembly against GenBank and Indigo genomes at an ANI ≥ 95% threshold. The known geographic distribution of *Tardiphaga zeae* sp. nov. in the United States is illustrated in Figure 10.

### Morphology, physiology and biochemical characteristics

#### *Paenibacillus glycinicola* sp. nov. strain SS118, and *P. triticicola* sp. nov. strain PI119

The two *Paenibacillus* strains, SS118 and PI119 had similar morphological characteristics but important differences were found. On R2A agar, colonies of SS118 were irregular and flat, with undulate margins, and were non-pigmented. The same colony characteristics were recorded for PI119 on Tryptic Soy agar (TSA), as this strain did not grow on R2A. Concerning Gram staining, SS118 was variable, while PI119 yielded a positive reaction. Cells of both strains are rod-shaped but differ in size, with SS118 measuring 2 – 4 µm long and those of PI119 only 1 – 2 µm (Figure 11A, Figure 11B). Motility was recorded for strain PI119 and showed growth at 30 and 37°C, while strain SS118 is non-motile and only grew at 30°C. Neither strain grew at 42°C. The new *Paenibacillus* strains are aerobic with a respiratory metabolism. Catalase production was detected in PI119, but neither strain produced oxidase nor urease. Physiological characterization using the Biolog GenIII system showed that the strains could use various common compounds as carbon and energy sources: D-maltose, D-cellobiose, lactose, melibiose, D-salicin, α-D-glucose, D-fructose, D-galactose and sodium lactate (1%).

**Figure 11.**
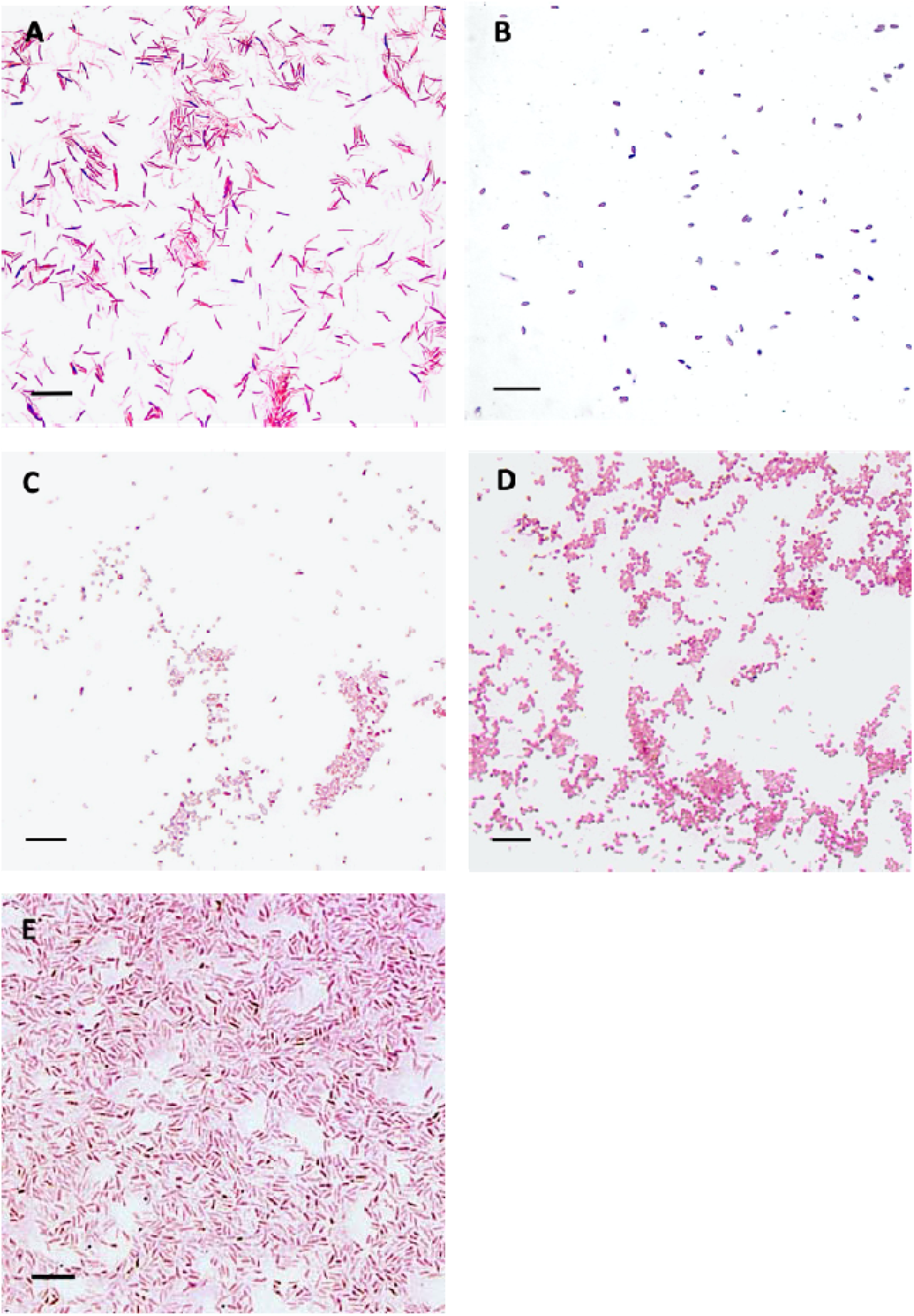
Morphology of new species depicted following Gram stain under bright field microscopy. A. *Paenibacillus glycinicola* sp. nov. strain SS118. B. *Paenibacillus triticicola* sp. nov. strain PI119. C. *Pseudescherichia oryzae* sp. nov. strain SS120. D. *Pseudomonas gossypii* sp. nov. PI121. E. *Tardiphaga zeae* sp. nov. strain SS122. Bar = 10 µm.

Differences were found, as only strain PI119 metabolized D-trehalose, sucrose, D-turanose, stachyose, D-raffinose, β-Methyl-D-glucoside, D-sorbitol D-mannitol and glycerol. Other results are given in the corresponding species descriptions. Concerning NaCl tolerance, both strains grew at concentrations up to 8%. In addition, the new bacteria were sensitive to the antibiotics troleandomycin, lincomycin and rifamycin SV, however, PI119 showed resistance to minocycline.

Distinguishing phenotypic characteristics between *Paenibacillus glycinicola* sp. nov. and the closest phylogenetic type strains *P. helianthi* P26E^T^ and *P. sonchi* X19-5^T^ is summarized in Table 2. Differences between *P. triticicola* sp. nov. and *P. peoriae* NBRC 1554^T^ and *P. kribbensis* AM49^T^ are given in Table 3.

**Table 2.**
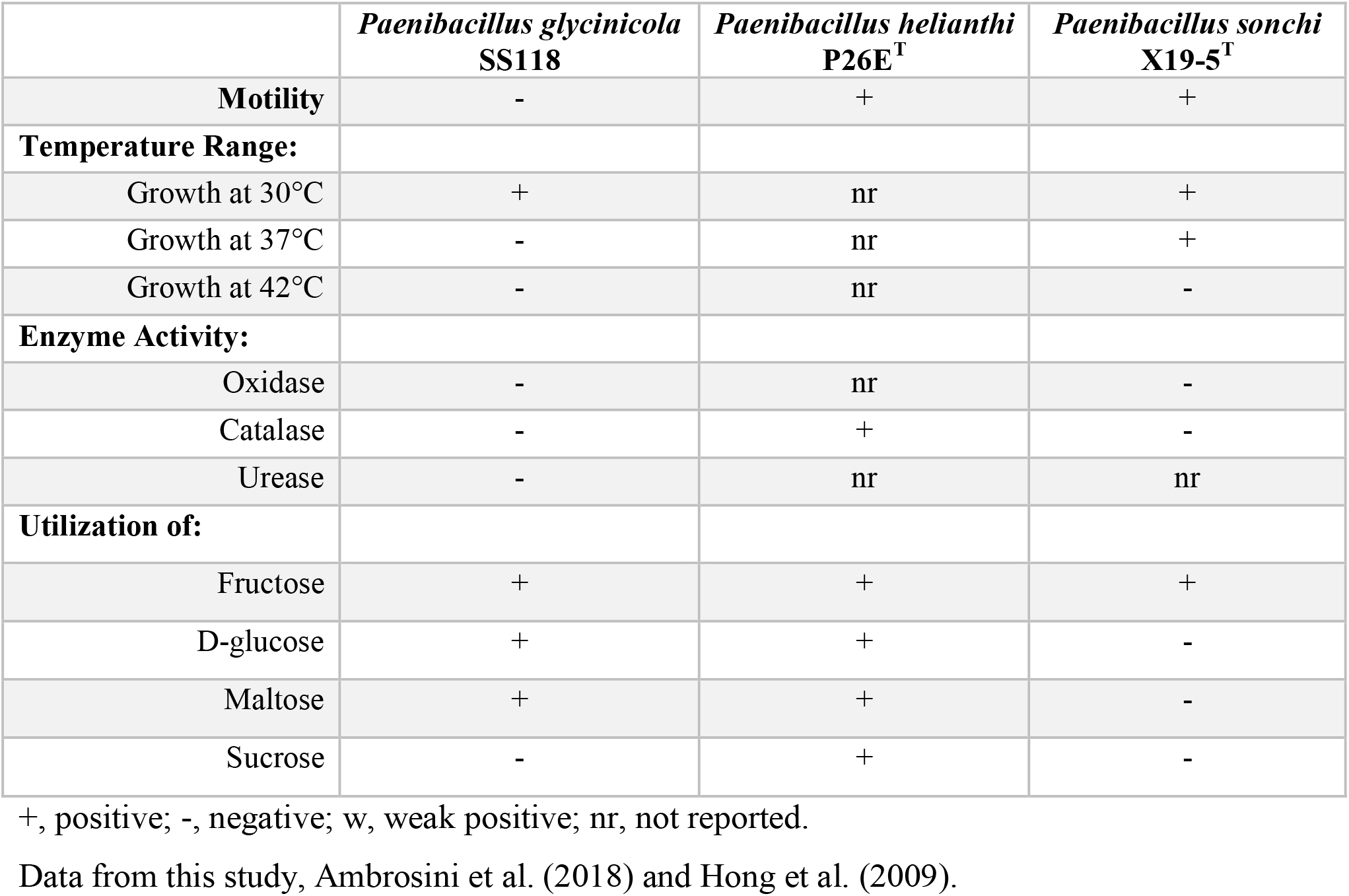
Distinguishing characteristics of *Paenibacillus glycinicola* sp. nov. strain SS118 and closely related species.

**Table 3.** Distinguishing characteristics of *Paenibacillus triticicola* sp. nov. strain PI119 and closely related species.

|  | *Paenibacillus triticicola*<br>PI119 | *Paenibacillus peoriae*<br>NBRC 15541<sup>T</sup> | *Paenibacillus*<br>*kribbensis* AM49<sup>T</sup> |
| --- | --- | --- | --- |
| **Temperature Range:** |  |  |  |
| Growth at 42°C | - | nr | nr |
| **Enzyme Activity:** |  |  |  |
| Oxidase | - | - | - |
| Catalase | + | + | + |
| Urease | - | nr | - |
| **Utilization of:** |  |  |  |
| Dextrin | + | - | - |
| N-Acetyl-D-Glucoside | - | + | + |
| L-Rhamnose | - | + | + |
| *myo*-Inositol | - | + | + |

+, positive; -, negative; w, weak positive; nr, not reported.
Data from this study, Montefusco et al. (1993), Wang et al. (2013), and Yoon et al. (2003).

#### *Pseudescherichia oryzae* sp. nov. strain SS120

The cells of strain SS120 are non-motile and stain Gram-negative. On TSA agar, circular, non-pigmented, and umbonate colonies with undulate margins are observed. Cells are small rod-shaped and measure 0.5 – 1µm long (Figure 11C). Strain SS120 can grow 30 – 42° and at pH 5 and 6, and in the presence of up to 4% NaCl. There is a weak reaction for catalase production but not for oxidase or urease. Under the Biolog GenIII system to assess carbohydrate utilization, the strain weakly catabolized D-maltose, D-trehalose, D-cellobiose and gentobiose. Other sugars used as carbon sources include glycerol, D-glucose-6-phosphate and D-fructose-6-phosphate. Other results are given in the species description. There is resistance to rifamycin SV and lincomycin, but not to minocycline. SS120 is also capable of growing in the presence of the chemicals tetrazolium blue and tetrazolium violet, but only shows weak growth against lithium chloride.

Differential phenotypic characteristics between strain SS120 and the only representative of the genus *Pseudescherichia, P. vulneris* NBRC 102420^T^ are presented in Table 4.

**Table 4.** Distinguishing characteristics of *Pseudescherichia oryzae* sp. nov. strain SS120 and closely related species.

|  | *Pseudescherichia oryzae*<br>SS120 | *Pseudescherichia*<br>*vulneris*<br>NBRC 102420<sup>T</sup> |
| --- | --- | --- |
| **Motility** | - | + |
| **Temperature Range:** |  |  |
| Growth at 30°C | + | nr |
| Growth at 37°C | + | nr |
| Growth at 42°C | + | nr |
| **Enzyme Activity:** |  |  |
| Catalase | w | + |
| **Utilization of:** |  |  |
| D-glucose | w | + |
| Lactose | w | + |
| Sucrose | - | + |
| D-mannitol | w | + |
| D-sorbitol | + | - |
| Raffinose | - | + |
| Glycerol | + | - |

+, positive; -, negative; w, weak positive; nr, not reported.
Data from this study and Brenner et al. (1982).

#### *Pseudomonas gossypii* sp. nov. strain PI121

Strain PI121 stained Gram-negative. On TSA, flat, yellow-pigmented colonies with entire margins are observed. Cells are rod-shaped and motile, measuring approximately 0.5 – 1 µm long (Figure 11D). Growth was recorded at 30 and 37°C but not at 42°C. The strain also grew at pH 5 and 6 and tolerated NaCl concentrations up to 4%. Strain PI121 had a weak catalase reaction. On the contrary, it did not produce oxidase or urease. Being aerobic, the strain has a respiratory metabolism. Based on the Biolog GenIII system, strain PI121 can use a diverse range of compounds as carbon and energy sources. These include D-trehalose, sucrose, α-D-glucose, D-mannose, D-fructose, D-galactose, D-sorbitol, D-mannitol, D-arabitol, myo-inositol and glycerol in addition to several organic acids. Other results are given in the species description. The strain shows resistance to rifamycin SV and lincomycin but is susceptible to minocycline, and nalidixic acid. The strain grows in the presence of several chemical inhibitors, which include tetrazolium violet, tetrazolium blue, lithium chloride, potassium tellurite, and Tween 40.

Distinguishing characteristics between the new strain and the closest phylogenetic type strains of the species, *P. luteola* NBRC 103146, *P. oryzihabitans* LMG 7040 and *P. oleovorans* IAM 1508, are given in Table 5.

**Table 5.** Distinguishing characteristics of *Pseudomonas gossypii* sp. nov. strain PI121 and closely related species.

|  | *P. gossypii*<br>PI121 | *P. oryzihabitans*<br>LMG7040 | *P. oleovorans*<br>IAM1508<sup>T</sup> | *P. luteola*<br>NBRC103146<sup>T</sup> |
| --- | --- | --- | --- | --- |
| **Temperature Range:** |  |  |  |  |
| Growth at 42°C | - | - | + | + |
| **NaCl tolerance:** |  |  |  |  |
| 8% | w | - | - | - |
| **Enzyme Activity:** |  |  |  |  |
| Oxidase | - | - | nr | - |
| Catalase | w | + | nr | + |
| Urease | - | + | nr | + |
| **Utilization of:** |  |  |  |  |
| Maltose | w | + | - | + |
| Gluconate | + | + | - | + |
| D-Sorbitol | + | + | - | - |
| Propionic Acid | w | - | - | nr |
| D-mannose | + | + | - | + |
| D-galactose | + | + | - | nr |
| Mannitol | + | + | - | + |

+, positive; -, negative; w, weak positive; nr, not reported.
Data from this study, Hauser et al. (2004), Liu et al. (2008), and Romanenko et al. (2005).

#### *Tardiphaga zeae* sp. nov. strain SS122

Cells of strain SS122 stain Gram-negative. On R2A agar colonies are non-pigmented, circular, flat with entire margins, and shows very slow growth similar to *Tardiphaga robiniae* LMG 26467^T^, the only representative of the genus. Cells are rod-shaped, motile and small, measuring less than 2 µm. The strain can grow at 30°C but not at 37 or 42°C, and at pH 6 but not pH 5. Carbon source characterization with the Biolog Gen III system showed that the strain can use fusidic acid, glucuronamide, quinic acid, α-keto-glutaric acid, D-malic acid, L-malic acid, and acetic acid. The strain shows resistance to aztreonam, nalidixic acid, lincomycin, minocycline and rifamycin SV.

Several phenotypic differences between strain SS122 and *Tardiphaga robiniae* LMG 26467^T^ are given in Table 6.

**Table 6.** Distinguishing characteristics of *Tardiphaga zeae* sp. nov. strain SS122 and closely related species.

|  | *Tardiphaga zeae*<br>SS122 | *Tardiphaga robiniae*<br>LMG 26467<sup>T</sup> | *Rhodopseudomonas*<br>*bookkerdii* NBRC<br>106595<sup>T</sup> |
| --- | --- | --- | --- |
| **Motility** | + | - | + |
| **Enzyme Activity:** |  |  |  |
| Catalase | - | + | + |
| **Utilization of:** |  |  |  |
| D-glucose | - | w | - |
| Malate | + | - | + |

+, positive; -, negative; w, weak positive; nr, not reported.
Data from this study and De Meyer et al. (2012).

### DESCRIPTION OF *PAENIBACILLUS GLYCINICOLA* SP. NOV

#### *Paenibacillus glycinicola* (gly.ci.ni’co.la. N.L. fem. n. *Glycine*, the genus of the soybean; L. masc./fem. n. suff. -cola, inhabitant dweller; from L. masc./fem. n. *incola*, dweller; N.L.)

Gram variable, rod-shaped, nonmotile bacterium (2-4 µm long). Irregular, non-pigmented colonies on R2A agar. Does not grow on TSA. Growth observed at 30°C but not at 37 or 42 °C. Grows at pH 6 but not at 5 and on 8% (w/v) NaCl. Neither catalase, oxidase nor urease are produced. Weakly uses gentiobiose as a carbon source, but not N-acetyl-D-glucosamine, N-acetyl-β-D-mannosamine, N-Acetyl-D-galactosamine, N-Acetyl-neuraminic acid, D-mannose, 3-methyl-glucose, DL-fucose, L-rhamnose or inosine. Most amino acids are not used as carbon sources. Growth is not inhibited by potassium tellurite.

The type strain SS118 was isolated from a stem of a healthy field-grown plant of *Glycine max* in Iowa.

The G+C content is 49.4%.

### DESCRIPTION OF *PAENIBACILLUS TRITICICOLA* SP. NOV

#### *Paenibacillus triticicola* (tri.ti.ci.co’la. L. neut. n. *Triticum*, wheat; L. masc./fem. n. suff. -*cola*, inhabitant dweller; from L. masc./fem. n. *incola*, dweller; N.L. masc./fem. n. *triticicola*, wheat dweller)

Gram stain positive, rod-shaped, motile bacterium (1 – 2 µm long). Irregular, non-pigmented colonies with undulate margin on TSA agar. Growth is observed at 30 and 37°C but not at 42°C. Grows at pH 6 but only weakly at pH 5 and in 4% (w/v) NaCl. Catalase is produced but not oxidase or urease. The following compounds are used as carbon sources: dextrin, D-trehalose, sucrose, D-turanose, stachyose, raffinose, β-methyl-D-glucoside, D-mannose, D-sorbitol, D-mannitol, glycerol, pectin, and D-gluconic acid. Growth not inhibited by tetrazolium violet and lithium chloride.

The type strain PI119 was isolated from the roots of *Triticum aestivum* collected in South Australia.

The G+C content is 45.6%.

### DESCRIPTION OF *PSEUDESCHERICHIA ORYZAE* SP. NOV

#### *Pseudescherichia oryzae* (o.ry’zae. L. gen. fem. n. *oryzae*, of rice)

Gram stain-negative, rod-shaped (0.5-1µm long), non-motile bacterium. Circular, non-pigmented, umbonate colonies with undulate margin on TSA agar. Growth observed at 30°C but not at 37 or 42°C. Grows at pH 6 but not at pH 5 and in 4% (w/v) NaCl. Catalase is weakly produced but not oxidase or urease. Dextrin, D-maltose, D-trehalose, D-cellobiose, gentiobiose, α-D-lactose, β-methyl-D-glucoside, D-salicin, D-galactose, inosine, 1% sodium lactate, fusidic acid, D-sorbitol, D-arabitol, glycerol, D-glucose-6-PO_4_ and D-fructose-6-PO_4_. Growth is not inhibited by tetrazolium violet, tetrazolium blue.

The type strain SS120 was isolated from a field-grown *Oryza sativa* seedling in Arkansas.

The G+C content is 55.9%.

### DESCRIPTION OF *PSEUDOMONAS GOSSYPII* SP. NOV

#### *Pseudomonas gossypii* (gos.sy’pi.i. N.L. gen. neut. n. *gossypii*, of the cotton genus *Gossypium*)

Gram stain-negative, rod-shaped (0.5 – 1µm long), motile bacterium. Circular, yellow-pigmented, colonies on TSA agar. Growth observed at 30 and 37°C but not at 42 °C. Grows at pH 5 and 6 and in 4% (w/v) NaCl. Catalase is weakly produced but not oxidase or urease. In addition to the substrates above, the following compounds are used as carbon sources: 1% sodium lactate, fusidic acid, inosine, D-sorbitol, D-arabitol, glycerol, D-glucose-6-PO4, D-fructose-6-PO4, DL-galacturonic acid, D-gluconic acid, D-glucuronic acid, mucic acid, D-saccharic acid, lactic acid and citric acid. Grows in the presence of tetrazolium violet and tetrazolium blue, niaproof 4, and weakly in potassium tellurite.

The type strain PI121 was isolated from a cotton bowl of a healthy field-grown *Gossypium hirsutum* in New Mexico.

The G+C content is 65.1%.

### DESCRIPTION OF *TARDIPHAGA ZEAE* SP. NOV

#### *Tardiphaga zeae* (ze’ae. L. gen. fem. n. *zeae*, of *Zea mays*)

Gram stain-negative. Rod-shaped (<2µm long), motile, and slow growing bacterium. Circular, non-pigmented colonies on R2A, does not grow on TSA agar. Growth observed at 30°C but not at 37 or 42 °C. Grows at pH 6 but not at pH 5. Does not grow in the presence of NaCl. Produces oxidase but not urease. Catalase is weakly produced. Based on the Biolog GENIII system only a few compounds are used as carbon sources: D-serine, 1% sodium lactate, fusidic acid, D-gluconic acid, glucuronamide, quinic acid, methyl pyruvate, L-lactic acid, acetic acid, and formic acid. The following sugars do not serve as carbon sources: dextrin, D-maltose, D-trehalose, D-cellobiose, D-turanose, gentiobiose, sucrose, stachyose, D-raffinose, α-D-lactose, D-melibiose, D-salicin, α-D-glucose, D-mannose, D-fructose, D-galactose, D-fucose, L-rhamnose, inosine, D-sorbitol, D-mannitol, D-arabitol, myo-inositol and glycerol. Gelatin is not hydrolyzed.

The type strain SS122 was isolated from a root of healthy field-grown *Zea mays* in Iowa. The G+C content is 61.4%.

## ACKNOWLEDGEMENTS

We thank Dr. Aljosa Trmcic, Wiedmann Lab, Cornell University, for pointing us toward the Food Microbe Tracker. We also thank Professor Aharon Oren, The Hebrew University of Jerusalem, for help with nomenclature.

